# Loss of TRIM21 drives UVB-induced systemic inflammation by regulating DNA-sensing pathways

**DOI:** 10.1101/2024.04.10.588897

**Authors:** Gantsetseg Tumurkhuu, Richard Moore, Graziela Perri, Lihong Huo, Arati Naveen Kumar, Gabriela de los Santos, David Gibb, Jessica Carriere, Jeong Min Yu, Rachel Abuav, Daniel J. Wallace, Mariko Ishimori, Wonwoo Shon, Andrea Dortfleutner, Christian Stehlik, Caroline A. Jefferies

## Abstract

**Background:** Exposure of systemic lupus erythematosus (SLE) patients to ultraviolet light B (UVB) triggers local and systemic inflammation, with cytosolic DNA sensing and induction of type I interferons (IFNs) known to play a role. We previously identified TRIM21 as a negative regulator of DNA sensing and IFN expression. <u>Here we explore the role of TRIM21 in regulating local and systemic responses following UVB exposure.</u>

**Methods:** WT (C57BL/6) and *Trim21^-/-^* mice were irradiated with UVB (100mJ/cm^2^) daily for 1 and 3 weeks, and UVB-induced inflammation in skin, blood, and spleen were analyzed by qPCR, histology, RNA sequencing and flow cytometry. Mechanistic studies were performed in bone marrow-derived macrophages (BMDMs) and mouse skin fibroblasts (MDF) from WT and *Trim21^-/-^* mice, and *TRIM21^-/-^*THP-1 cells.

**Results:** Infiltration of inflammatory cells and induction of type I IFN developed in UVB-exposed areas in both sets of mice, however *Trim21^-/-^*mice developed splenomegaly, enhanced total IgG levels and IFN-stimulated genes (ISG) in the blood and spleen. Enhanced basal and UVB-dependent *Ifnb1* expression was observed in *Trim21^-/-^* BMDMs and MDFs, which was dependent on the cytosolic DNA sensing cGAS-STING pathway. Mechanistically, we found both degradation of DDX41 and STING levels were impaired in stimulated *Trim21^-/-^*BMDMs.

**Conclusion:** Taken together, our results indicate that TRIM21 protects against IFN induction at local and systemic levels through restricting STING signaling. Our finding that reduced levels of TRIM21 are observed in SLE patients with cutaneous involvement indicates a potential role for TRIM21 in guarding against systemic flare in SLE patients.

## Introduction

Abnormal responses to ultraviolet B (UVB) light or photosensitivity are experienced by approximately two thirds of systemic lupus erythematosus (SLE) patients, with sun-exposure triggering either local skin inflammation, increased systemic activity (flare), or both (1). The mechanisms that predispose patients to develop systemic disease flares in response to sun exposure are unclear and are an important gap in our knowledge regarding the pathophysiology of this disease. Elevated type I interferon (IFNα, β and κ) and IFN-stimulated genes (ISG) are the most common immunologic abnormality in SLE and associate with disease severity and organ involvement, including cutaneous involvement (2–5). Photosensitivity to UVB is a well-known trigger of both localized cutaneous disease (CLE) and systemic disease. In cutaneous manifestations of SLE, high levels of type I IFNs are detected in both lesional and non-lesional skin and correlate with systemic ISG expression (6). UVB exposure has multiple effects on keratinocytes, the first line of defense against exposure, including IFN induction (2, 7). However, the molecular links between UVB-photosensitivity and development of systemic disease are unknown.

Mechanistically, UVB exposure triggers several biologic pathways in the skin including DNA damage, DNA repair, cell death, and release of self-nucleic acids. Self-RNA and DNA (mitochondrial and nucleic) act as potent damage-associated molecular patterns (DAMPs), and are detected by cytosolic nucleic acid sensors (5, 8–12) such as cGAS and DDX41 (13–15). cGAS, once it binds dsDNA, synthesizes the cyclic dinucleotide cyclic-GMP-AMP (cGAMP) which acts as a second messenger in the cell, binding and activating the adaptor protein STING to drive IFNβ induction (16–19). DDX41 can also bind cGAMP and STING and drive IFNβ expression (13, 14). Recent work has clearly implicated cGAS and STING in UVB-mediated IFN expression in the skin of C57BL/6 mice, with the absence of STING or cGAS preventing IFN induction in the skin following UVB exposure (20, 21).

TRIM21, otherwise known as Ro52, is a common autoantigen associated with cutaneous SLE and is commonly upregulated in photo-provoked skin (7). Moreover, anti-Ro52/TRIM21 antibodies accumulate in CLE skin and exacerbate local inflammatory responses. As an E3 ubiquitin ligase, TRIM21 negatively regulates type I IFN induction downstream of RNA and DNA sensing pathways by ubiquitinating and targeting DDX41 and the transcription factors IRF3, 5 and 7 for proteasomal degradation (22–30). In doing so, TRIM21 limits the IFN response and prevents autoimmune disease. *Trim21*-deficient mice develop splenomegaly and a lupus-like systemic disease as a result of enhanced TLR7 and 9-induced IFN induction, driven by increased levels of IRF3, 5 and 7 (27). Importantly *Trim21^-/-^* mice exhibit skin lesions that are provoked by DNA damage, suggesting *Trim21* may play an important role in skin-involvement in SLE (27).

In this study we explored the potential that loss of *Trim21* might exacerbate UVB-induced skin and systemic inflammation. We found *Trim21-*deficiency results in enhanced local and systemic inflammation and ISG expression of UVB-exposed mice. In addition to increased DDX41 stability, we observed *Trim21^-/-^* cells fail to degrade STING after engaging the cGAS/STING pathway, resulting in upregulated type I IFN and ISGs. Our findings show that TRIM21 is an important regulator of systemic IFN responses following UVB challenge in the skin and suggest that loss of TRIM21 expression or activity in the skin of SLE patients with cutaneous involvement may enhance UVB-induced systemic involvement.

## Materials and Methods

### Mice

*Trim21^GFP/GFP^* mice on C57BL/6 background (*Trim21^-/-^*) were a gift from the Dr. Wahren-Herlenius lab (27). C57BL/6 and *Sting^gt/gt^*mice (on C57BL/6 background) were purchased from The Jackson Laboratory. Mice were propagated and maintained in the animal facility at the Department of Comparative Medicine (Davis Building) of Cedars-Sinai Medical Center. Female animals only were used in this study and were between 8-12 weeks old at the time of experimentation. All experiments procedure was approved by institutional IACUC committee.

### UVB irradiation

Eight-week-old wild-type (WT) and *Trim21^-/-^* were shaved dorsally prior to UVB exposure. Mice were allowed to move freely in their cage during UVB exposure and mice were treated with/without UVB 100 mJ/cm^2^ daily for one or three weeks.

### Isolation of skin, spleen cells and flow cytometry

Skin lesion, spleen, and blood were harvested following final UVB exposure. To generate a single cell suspension, spleens were passed through a 70-μM filter. Skin samples were minced and digested with 0.28 units/ml Liberase TM (Roche) and 0.1 mg/ml of Deoxyribonuclease I (Worthington) for 60 minutes at 37°C, then passed through a 70-μM filter and washed with 1% bovine serum albumin in phosphate buffered saline. Cells were then incubated in flow block (1% bovine serum albumin and 1% horse serum in PBS) for 30 min, followed by staining with mouse-specific fluorescent antibodies CD45 Alexa Fluor 700 (clone:30-F11), CD3 PE/Cy7 (clone 145-2C11), CD4 APC/Cy7 (clone GK1.5), CD11b Pacific Blue (clone M1/70), Ly6G APC (clone 1A8), CCR2 PE (clone SA203G11), all purchased from BioLegend (San Diego, CA); Siglec1 BV421 (clone 7-239) (BD Bioscience) for 45 minutes. All samples were processed using SA3800 flow cytometer (Sony) and data analyzed with FlowJo VX.0.7 software (Tree Star).

### Gene expression analysis

RNA was isolated from full-thickness skin samples using RNeasy Fibrous Tissue Mini Kit with on-column DNase treatment (Qiagen). From whole blood, splenocytes, and in vitro cultured cells, RNA was isolated using TRI Reagent (Millipore Sigma). First-strand complementary DNA (cDNA) was generated using 500 ng RNA with a High-Capacity cDNA Reverse Transcription Kit using random primers (Applied Biosystems). Reactions were run in triplicate on a StepOnePlus real-time PCR instrument (Applied Biosystems) using gene-specific primers. Ct values were standardized to the house-keeping gene *Rn18s* (mouse) or *RNA18SN* (human).

The following murine primer sequences were used:

*Rn18s*, 5′-GAGGGAGCCTGAGAAACGG-3′ (F) 5′- GTCGGGAGTGGGTAATTTGC-3′ (R);
*Cxcl10*, 5′-GCCGTCATTTTCTGCCTCAT-3′ (F) 5′- GCTTCCCTATGGCCCTCATT-3′ (R);
*Ifnb1*, 5’-CAGCTCCAAGAAAGGACGAAC-3’(F) 5’- GGCAGTGTAACTCTTCTGCAT-3’ (R);
*Isg15*, 5’-GGTGTCCGTGACTAACTCCAT-3’ (F) 5’- TGGAAAGGGTAAGACCGTCCT-3’ (R).

The following human primer sequences were used:

*IFNB1*,5’-CTTGGATTCCTACAAAGAAGCAGC-3’(F);5’- TCCTCCTTCTGGAACTGCTGCA-3’(R)

### Analysis of anti-IgG and dsDNA IgG antibody serum levels

Serum was collected post UVB treatment. Total IgG and anti-dsDNA IgG antibody levels were analyzed via ELISA kits (6320 and 5120, Alpha Diagnostic, San Antonio, TX).

### Histopathology, immunohistochemistry, and immunofluorescence

Tissue samples were immersed in 10% buffered formalin for 48 hours at 4°C, paraffin embedded, and stained using hematoxylin and eosin. For immunofluorescence, following the addition of secondary antibodies, slides were stained with DAPI, washed, air-dried, and mounted. Images were taken on a Zeiss Axio Observer and anti-IgG intensity was quantified using ImageJ.

### Patients and Healthy Controls

This study was approved by the institutional review board at Cedars-Sinai Medical Center and in line with the principles of the Declaration of Helsinki. All participants provided informed consent before joining. Participants were recruited from the lupus clinic. Healthy controls were age matched and recruited from Cedars-Sinai Medical Center. Blood samples were collected from all patients and controls using PAXgene blood RNA tubes.

### PAXgene RNA isolation

Blood samples were kept in PAXgene RNA tubes in -80 until ready to process. RNA was isolated using the PAXgene Blood RNA kit according to the manufacturers guidelines (PreAnalytiX). Eluted RNA was dissolved in RNase-free water. The quality and quantity of RNA were evaluated using the Agilent 2100 BioAnalyzer (Santa Clara, CA, USA).

### Isolation, culture, and stimulation of cells and tissue

Bone marrow-derived macrophages (BMDMs) were cultured as previously described (31). Briefly, bone marrow was flushed from the tibia and femurs of 8-10 week old mice and differentiated for 7 days in DMEM supplemented with 10% fetal bovine serum (FBS, Omega Scientific Inc), 1X Gibco Antibiotic-Antimycotic (A/A, 15240062, Thermo Fisher Scientific), and 20% L929-conditioned media in a 37°C, 5% CO_2_ incubator.

Murine dermal fibroblasts (MDFs) and skin explants were cultured as previously described (32). Briefly, mice were euthanized and shaved dorsal skin was excised and digested with collagenase D and pronase. MDFs were cultured in RPMI-1640 Medium (11875119, ThermoFisher Scientific) supplemented with 10% FBS and 1X A/A. Skin explant assay was performed as previously described (33). Briefly, explants were excised from shaved dorsal skin of euthanized mice using 6-mm punch biopsies and cultured similarly to MDFs. Before stimulation, MDFs and skin explants were starved in 2% FBS-supplemented medium for 24 hours before exposure to 500 mJ/cm^2^ UVB.

2’-3-cyclic GMP-AMP (cGAMP) (tlrl-nacga23-1, InvivoGen) was added at 10μg/mL with or without 4μM H151 (inh-h151, InvivoGen) to cell supernatant (added 30 minutes prior to stimulation) for 1, 2, 4, or 8 hours. 50nM immunostimulatory dsDNA (ISD) was transfected into BMDMs or THP-1 cells with Lipofectamine (L3000008, ThermoFisher Scientific) for 1, 2, 4, or 8 hours. ISD was formed by annealing the following sequences: ISD Forward: TACAGATCTACTAGTGATCTATGACTGATCTGTACATGATCTACA ISD Reverse: TGTAGATCATGTACAGATCAGTCATAGATCACTAGTAGATCTGTA

### RNA Sequencing

RNA from splenocytes isolated from UVB exposed WT and *Trim21^-/-^*mice were analyzed for RNA integrity on the 2100 Bioanalyzer using the Agilent RNA 6000 Nano Kit (Agilent Technologies, Santa Clara, CA) and quantified using the Qubit RNA HS Assay Kit (ThermoFisher Scientific, Waltham, MA). RNA libraries were sequenced on a NovaSeq 6000 (Illumina, San Diego, CA) using 75bp single-end sequencing. On average, approximately 50 million reads were generated from each sample.

### Bioinformatics and data analysis

Raw reads obtained from RNA-Seq were aligned to the transcriptome using STAR (version 2.5.0/RSEM (version 1.2.25) with default parameters, using GRCm38 transcriptome reference downloaded from https://www.gencodegenes.org, containing all protein coding and long non-coding RNA genes based on human GENCODE version 33 annotation. Expression counts for each gene in all samples were normalized by a modified trimmed mean of the M-values normalization method and the unsupervised principal component analysis (PCA) was performed with DESeq2 Bioconductor package version 1.42.0 in R version 4.3. Each gene was fitted into a negative binomial generalized linear model, and the Wald test was applied to assess the differential expressions between two sample groups by DESeq2. Benjamini and Hochberg procedure was applied to adjust for multiple hypothesis testing, and differential expression gene candidates were selected with a false discovery rate less than 0.05. For functional enrichment analysis across sample groups, we conducted genes enrichment analysis using the R package “clusterProfiler v4.10.0”[10] and “pathfindR v2.3.1[11].

### Generation of *TRIM21^-/-^* and THP-1 cells and *TRIM21* knock-in of *TRIM21^-/-^* THP-1 cells

*TRIM21^-/-^* THP-1 cells were generated using CRISPR/Cas9 targeting. Two guide RNAs (gRNAs) were designed to target hTRIM21 Exon 2: gRNA#1: ATGCTCACAGGCTCCACGAA (Exon2 ORF bp67-86); gRNA#2: ATGTTGGCTAGCTGTCGATT (Exon2 ORF bp196-215) and cloned into lentiCRISPRv1 (Addgene, plasmid 49535). After lentiviral transduction using lentiCRISPRv1 as control as described below, clonal populations were screened by PCR amplification and sequencing of the targeted genomic sequence using the following primers:

hTrim21gRNA1-Fwd: GTCTCCACACTGCTGTTTAACG;
hTrim21gRNA2-Rev: TTCCCATCTTTCTCACAGAACA.

To restore *TRIM21* expression into *TRIM21^-/-^* THP-1 cells, hTRIM21 full-length CDS with N-terminal Myc and EGFP was cloned into pLEX-MCS and transfected into HEK293T-Lenti cells (Clontech 632180) along with pMD.2G and psPAX2 (Addgene, plasmids 12259 and 12260) to generate lentiviral particles. Forty-eight hours post-transfection, lentiviral suspensions were harvested, passed through 0.45μm filters and mixed with 106 TRIM21KO THP-1 cells in the presence of 0.5μg/mL polybrene. Cells were centrifuged at 1,800rpm for 50min at 32°C. After overnight incubation at 37°C, the medium was changed and EGFP^+^ cells were sorted 72h post-infection and expression of Trim21 was verified by western blot and cells were selected to match expression of endogenous Trim21.

### Western Blot

Cells were lysed after ISD transfection or cGAMP stimulation using RIPA lysis buffer (0.05% SDS, 0.5% NP-40, and 0.25% Na-Deoxycholate in PBS) and quantified using BCA Protein Assay Kit (ThermoFisher Scientific). 20μg sample per lane were loaded onto 4-20% precast polyacrylamide gel membrane and then transferred onto nitrocellulose membranes (BioRad). Membranes were labeled with anti-DDX41 (Novus Bio), anti-IRF3 (clone D83B9, Cell Signaling Technology), anti-STING (Novus Bio), anti-TRIM21 (clone EPR20290, abcam), and anti-β-actin (Cell Signaling Technology). Following incubation with HRP-conjugated anti-mouse or anti-rabbit IgG (Cell Signaling Technology), signals were visualized with SuperSignal West Pico PUS Chemiluminescent Substrate (ThermoFisher Scientific) on a ChemiDoc XRS+ Scanner (BioRad).

### STATISTICS

All data was graphed, and statistics performed using GraphPad Prism. For data comparing multiple groups, two-Way ANOVA with Fisher’s LSD test was used. Comparison between two groups was completed via a two-tailed Student’s t-test for normally distributed data. Comparisons were considered significant with a p value of <0.05.

## Results

### The loss of TRIM21 accelerates UVB induced cutaneous inflammation

Ultraviolet B (UVB) and C (UVC) light drive type I IFN in keratinocytes, and both acute and repeated UVB exposure result in upregulation of type I IFN and IFN-stimulated genes (ISGs) within the skin of mice (34). Mice lacking TRIM21 develop severe dermatitis and systemic autoimmunity in response to chronic physical injury (27). To test whether TRIM21 plays a role in regulating the IFN-response to chronic UVB exposure, we shaved *Trim21^+/+^*(WT) and *Trim21^-/-^* (KO) C57BL/6 mice and exposed them to a low dose of UVB light on consecutive days over three weeks. While both WT and *Trim21^-/-^* animals demonstrated similar levels of infiltrating CD45^+^ immune cells into the skin following chronic UVB exposure (Supplementary Figure 1), CD11b^+^ cells were increased in UVB-exposed *Trim21^-/-^* skin (Figure 1A) and had higher levels of the IFN-stimulated gene (ISG), Siglec1, compared to WT CD11b^+^ cells (Figure 1B). In addition, IFNβ and the IFN-stimulated gene *Cxcl10* were significantly upregulated in *Trim21-* deficient skin tissue compared to WT (Figure 1C), with skin from *Trim21-*deficient animals showing increased IgG in the skin after UVB compared to WT (Figure 1D).

**Figure 1.**
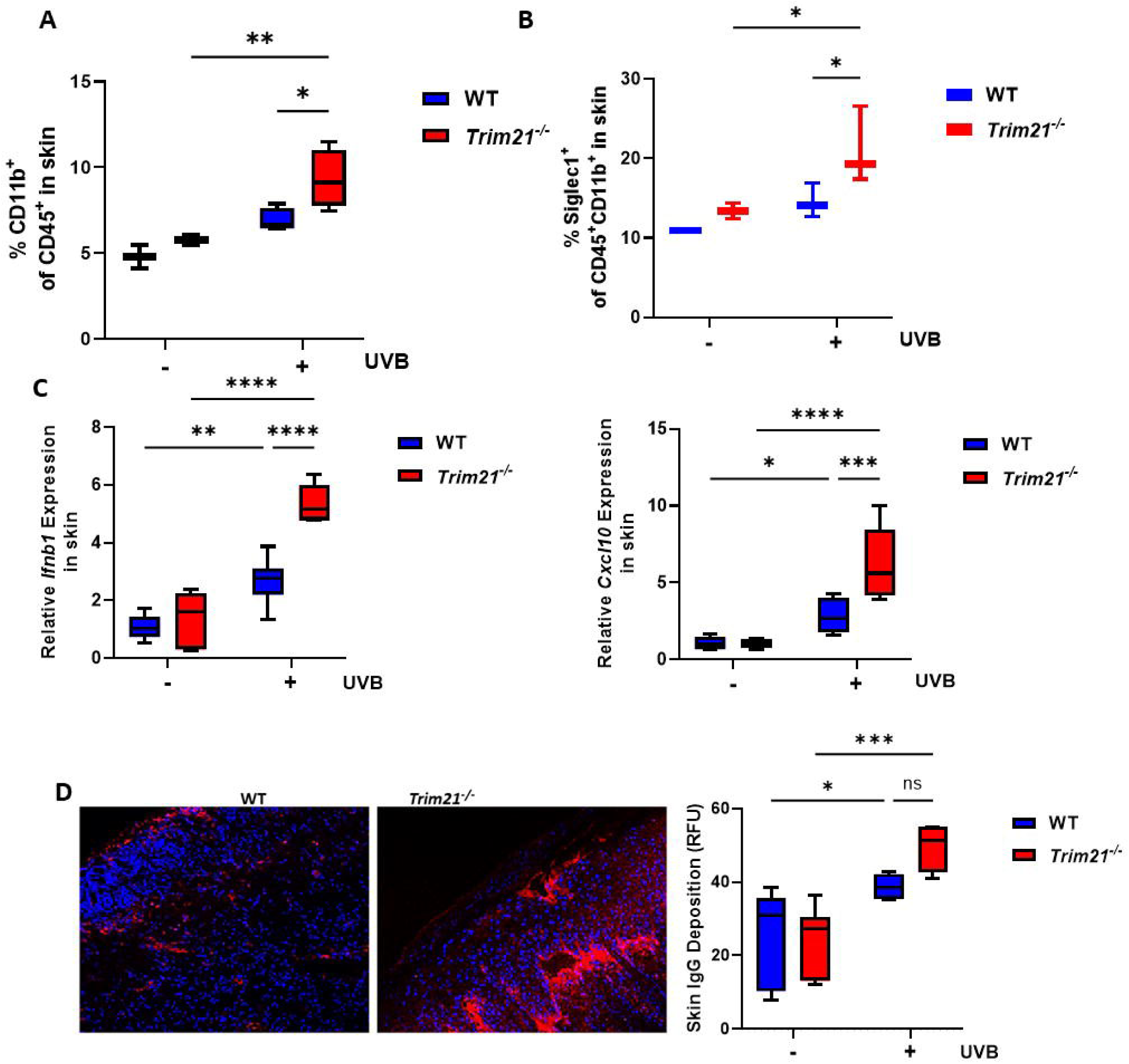
Ultraviolet B (UVB) skin irradiation induces a mixed local inflammatory response and higher cutaneous inflammation in *Trim21^-/-^* mice. . Flow cytometry analysis following the final dose of the 3-week UVB treatment showing the percentage of infiltrating **(A)** total myeloid cells in the skin and **(B)** Siglec1^+^ myeloid cells (CD11b^+^CD45^+^). **(C)** Cutaneous expression of *Ifnb1* and *Cxcl10* following 3-week UVB irradiation as determined by qPCR. **(D)** IgG deposition in UVB induced skin lesion with representative pictures and quantification analysis. All data represent mean ± SD. Statistical significance was determined using Two-Way ANOVA with Fisher’s LSD test; ns=no significance, *p<0.05, **p<0.01, ***p<0.001, and ****p<0.0001. In all cases data representing WT and Trim21^-/-^ mice is shown in blue and red bars, respectively.

### Loss of *Trim21* enhances the Systemic Inflammatory Response to UVB

In addition to the local immune response, acute UVB exposure also induces inflammation of the spleen and kidney (35). Because of this, we asked if TRIM21 protected against systemic inflammation due to chronic UVB exposure. We observed enhanced splenomegaly in *Trim21^-/-^* mice compared to WT (Figure 2A) with higher total levels of CD11b^+^ cells in the spleens of *Trim21^-/-^* mice (Supplemental Figure 1B). In addition, whilst splenic levels of CD11b^+^Ly6G^+^ cells increased in spleens of all UVB-exposed mice, the expansion was significantly higher in spleens of UVB-exposed *Trim21^-/-^* mice compared to WT (Figure 2B). In keeping with enhanced IFN responses, CD11b^+^ splenic populations from *Trim21^-/-^* mice had increased expression of the IFN-stimulated gene (ISG) Siglec1 compared to WT mice (Figure 2C). In addition, expression of *Ifnb1* and *Mx1* were enhanced in the spleens of UVB exposed *Trim21^-/-^* mice (Figure 2D). Noticeably, the percentage of CD4^+^ T cells in the spleen increased in the absence of *Trim21* (Figure 2E), with central memory CD4 T cells being more highly represented in the spleens of *Trim21^-/-^*mice compared to WT (Figure 2F). Pathway analysis of differentially expressed genes (represented in the volcano plot in Supplementary Figure 2A) identified increased expression of genes associated with mitosis and cell cycle regulation, DNA repair and ubiquitin ligase complexes (Supplementary Figure 2B), whereas the expression of genes associated with T cell activation and signaling through the IL2 receptor was decreased (data not shown), in keeping with our findings that central memory T cells were more highly represented in *Trim21^-/-^* splenocytes. Network analysis represents the increased enrichment of genes associated with DNA-damage and UV responses, and neutrophil activation and myeloid differentiation (Supplemental Figure 2C). Thus, loss of *Trim21* in mice results in systemic immune activation and enhanced neutrophil responses, ISG expression and activation of DNA damage and UV responsiveness in in response to UVB exposure of skin.

**Figure 2.**
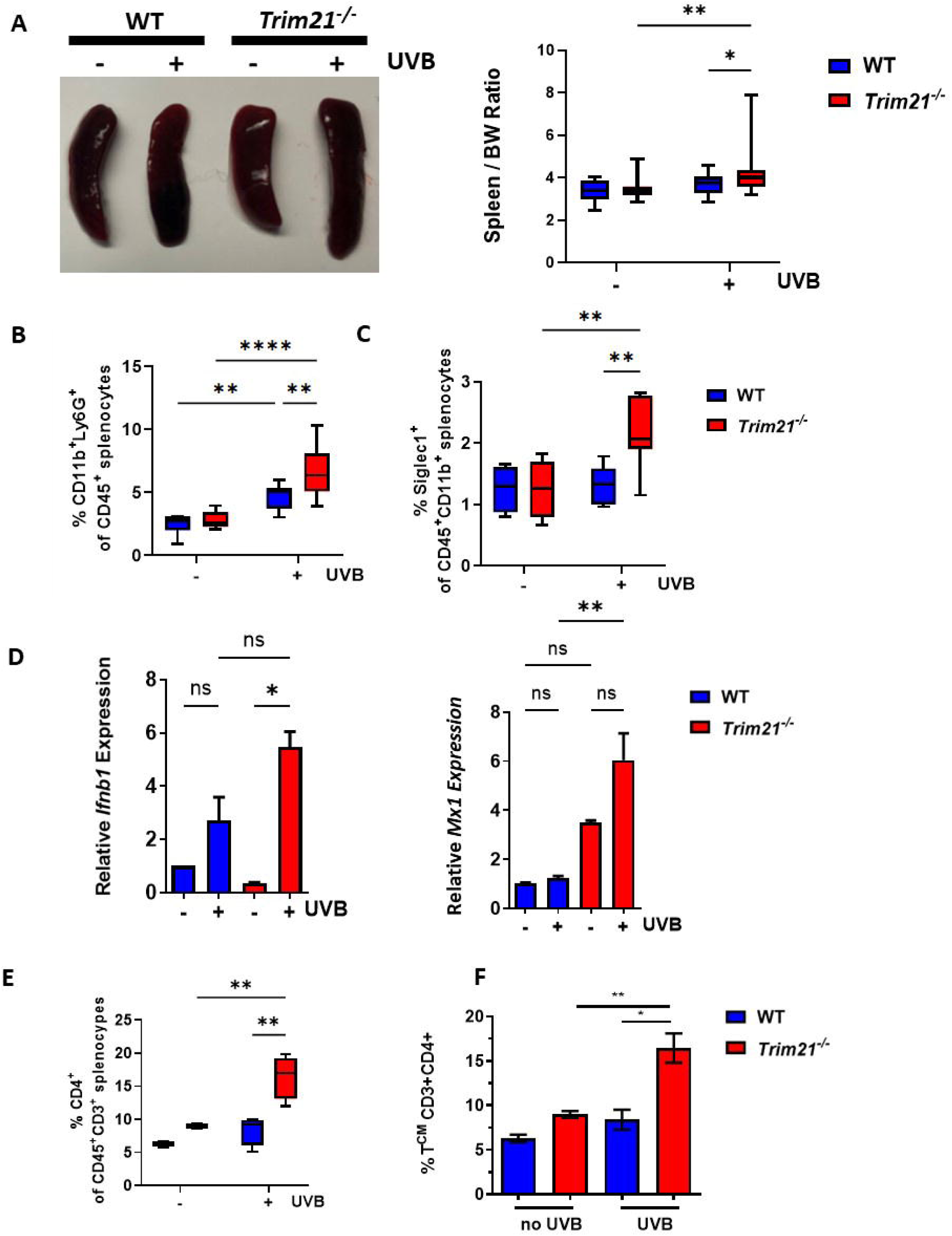
Splenic inflammation induced by chronic UVB irradiation is restrained by TRIM21. Following the final dose of the 3-week UVB treatment, **(A)** splenomegaly was measured by weight. Percentages of splenic **(B)** neutrophil cells and **(C)** Siglec1^+^ CD11b^+^ cells were analyzed via flow cytometry. **(D)** Splenocyte expression of *Ifnb1* and *Mx1* following 3-week UVB irradiation as determined by qPCR. Percentages of splenic **(E)** CD4^+^ T cells and **(F)** central memory T cells were analyzed via flow cytometry. All data represent mean ± SD and are representative of 3 independent experiments. Statistical significance was determined using Two-Way ANOVA with Fisher’s LSD test. *p<0.05, **p<0.01, ***p<0.001, and ****p<0.0001. In all cases data representing WT and Trim21^-/-^ mice is shown in blue and red bars, respectively.

Circulating CCR2^+^ monocytes are upregulated in response to UVB exposure and migrate to the UVB-exposed tissue to drive inflammation (21). In line with this, we observed increased CD11b^+^Ly6G^+^ circulating neutrophils and CD11b^+^Ly6C^+^ inflammatory monocytes in UVB-exposed *Trim21^-/-^* mice compared to WT controls (Figure 3A). Importantly, circulating CCR2^+^CD11b^+^ myeloid cells were increased compared to WT and expressed higher levels of Siglec1 (Figure 3B). In addition, *Trim21*^-/-^ mice had higher circulating IgG compared to WT (Figure 3C), although no increase in anti-dsDNA IgG was observed following three weeks of UVB exposure (data not shown). Altogether, these data support that loss of *Trim21* exacerbates the systemic inflammatory response to UVB exposure.

**Figure 3.**
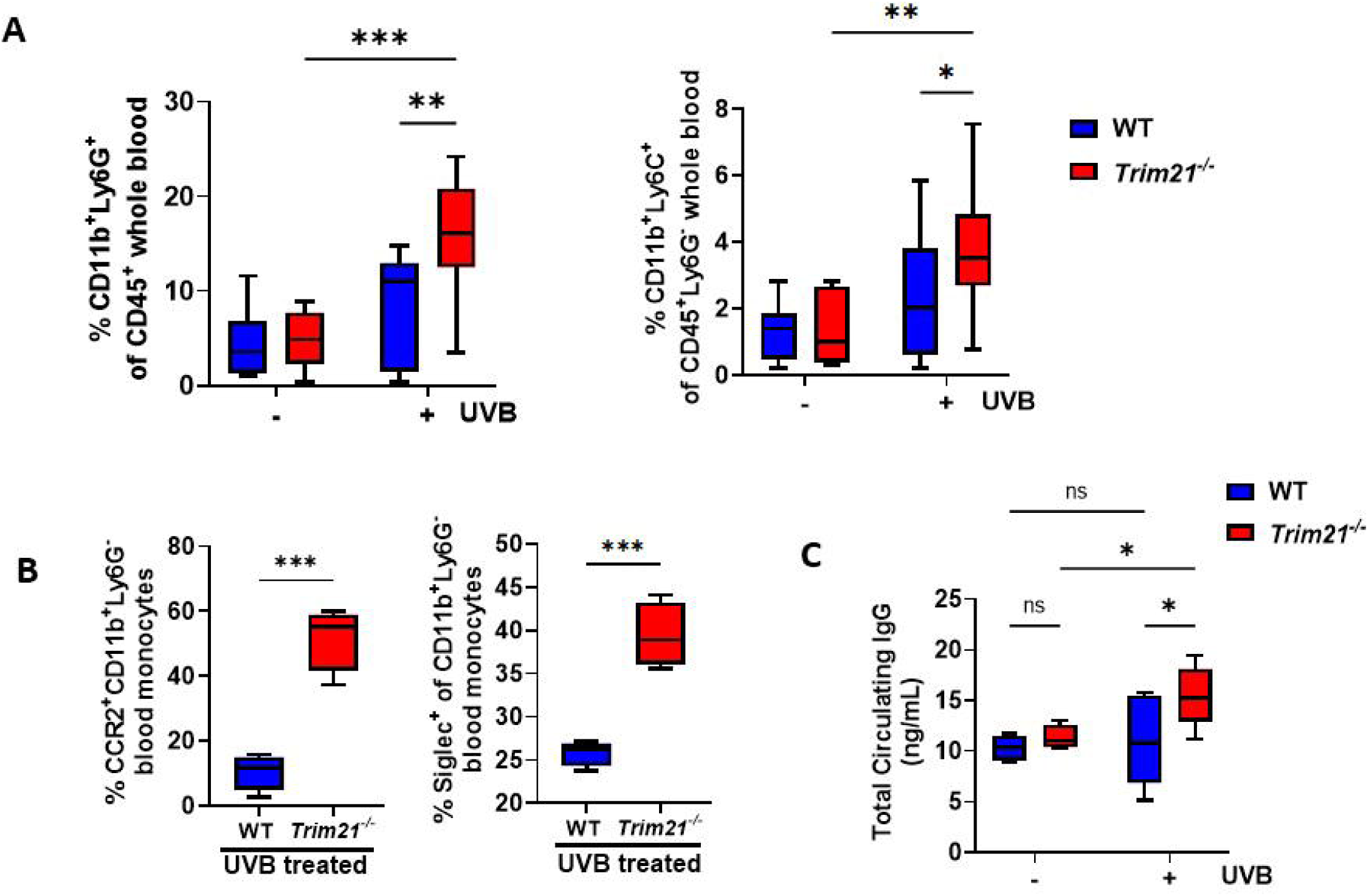
TRIM21-deficiency enhances systemic inflammation following UVB irradiation of the skin. Percentage of (A) neutrophils and inflammatory monocytes and of (B) CCR2^+^ and Siglec1^+^ CD11b^+^Ly6G^-^ cells from whole blood were analyzed via flow cytometry after the final dose of the 3-week UVB treatment. (C) Total circulating IgG was calculated via ELISA. All data represent mean ± SD and are representative of 3 independent experiments. Statistical significance was determined using Two-Way ANOVA with Fisher’s LSD test or unpaired t-test. ns=no significance, *p<0.05, **p<0.01, ***p<0.001. In all cases data representing WT and Trim21^-/-^ mice is shown in blue and red bars, respectively.

### TRIM21 Restricts the IFN-I Response via STING Signaling

The cGAS-STING pathway has recently been shown to regulate UVB-induced systemic responses in mice (21, 35, 36). In addition, the DNA sensor DDX41 is a target for TRIM21-mediated ubiquitination and degradation. To assess the effect of TRIM21-deficiency on DDX41 and STING stability, we transfected WT and *Trim21^-/-^* bone marrow-derived macrophages (BMDMs) with immunostimulatory dsDNA (ISD) to activate cGAS upstream of STING, then analyzed DDX41 and STING stability by western blotting. As previously published, DDX41 and STING were degraded in a time-dependent manner following ISD transfection of WT BMDMs, whereas both DDX41 and STING were stabilized in *Trim21^-/-^* BMDMs (Figure 4A). This was also recapitulated in cGAMP-stimulated THP1 cells in which *TRIM21* had been knocked out via CRISPR-Cas9, compared against *TRIM21*-deficient THP-1 cells stably reconstituted with *Myc-EGFP-TRIM21* (Supplemental Figure 3A, B). As expected, the increased stability of DDX41 and STING observed in the *Trim21^-/-^* BMDMs and *TRIM21*-deficient THP1 cells was mirrored by enhanced *Ifnb1/IFN*β*1* expression following activation of the cGAS-STING pathway (Figure 4B, red bars and Supplemental Figure 3C, grey bars, respectively). Importantly, functional deletion of *Sting* by crossing *Trim21^-/-^* with *Sting Golden ticket (Gt)* mice, which harbor a single nucleotide variant (T596A) of *Sting* that functions as a null allele (*Trim21^-/-^Sting^gt/gt^*BMDMs), reversed the enhanced *Ifnb1* expression observed in the *Trim21^-/-^*BMDMs {Sauer, 2011 #5537}(Figure 4B, grey bars). Similarly, loss of STING function in *Trim21^-/-^* BMDMs (either by genetic deletion using *Trim21^-/-^Sting^gt/gt^* BMDMs or using H151, a specific STING inhibitor (37)) potently reversed cGAMP-induced *Ifn*β*1* expression in *Trim21^-/-^*deficient BMDMs (Supplemental 4A&B, respectively). We next asked whether the increased type I IFN response in the skin of UVB-exposed *Trim21^-/-^*mice was dependent on STING signaling. Primary murine dermal fibroblasts (MDFs) from WT or *Trim21^-/-^*mice were UVB-treated with or without H151 (37), and assessed for *Ifnb1* expression by qPCR. As expected, UVB-treated *Trim21^-/-^* MDFs showed enhanced STING-dependent *Ifnb1* expression compared to WT MDFs (Figure 4C). Similarly, UVB-irradiated skin explants from WT, *Trim21^-/-^*and *Trim21^-/-^Sting^gt/gt^* mice demonstrated that *Trim21^-/-^* explants exhibited a STING-dependent increase of *Ifnb1* and *Isg15* expression (Figure 4D, Supplemental Figure 4C).

**Figure 4.**
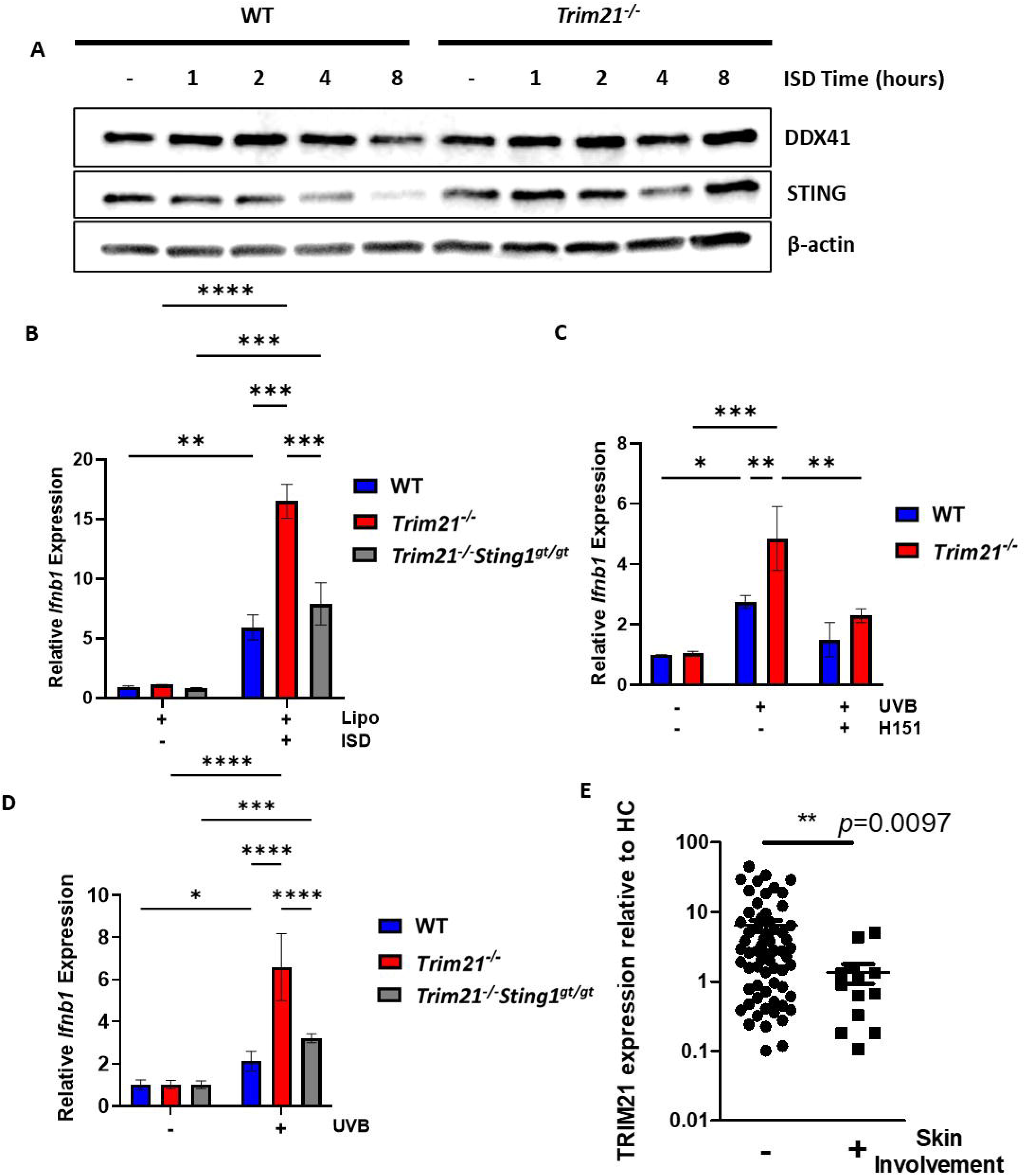
TRIM21 negatively regulates STING-dependent inflammation in response to UVB and cGAS-STING activation. **(A)** Bone marrow-derived macrophages (BMDMs) from WT or *Trim21^-/-^* mice were transfected with ISD for the designated time and protein levels of DDX41, STING, and β-actin were visualized via western blotting. Images shown are representative of n=3. **(B)** WT, *Trim21^-/-^*, and *Trim21^-/-^Sting^gt/gt^* BMDMs were transfected with ISD for 2 hours; (C) Murine dermal fibroblasts were isolated from WT and *Trim21^-/-^*mice and UVB irradiated (500mJ/cm2) and incubated for 24; **(D)** Whole WT, *Trim21^-/-^* or *Trim21^-/^Sting1^-/-^*skin explants were UVB-irradiated as for (C) and incubated for 24 hours; **(B-E)** *Ifnb1* and *Isg15* transcript levels were determined via qPCR. All transcript values are relative to unstimulated or transfection reagent-only treated (Lipo) cells. All data represent mean ± SD and are representative of 3 independent experiments. Statistical significance was determined using Two-Way ANOVA with Fisher’s LSD test. ns=no significance, *p<0.05, **p<0.01, ***p<0.001, and ****p<0.0001. In all cases data representing WT and Trim21^-/-^ mice is shown in blue and red bars, and *Trim21^-/^Sting1^-/-^*shown in grey.

Finally, given the protective effect of TRIM21 on UVB-driven responses in mice, we asked whether there was any alteration in *TRIM21* expression in SLE patients in our repository with documented cutaneous involvement at the time of blood draw. *TRIM21* levels were significantly lower in in whole blood samples from patients with cutaneous involvement (Figure 4E). Thus, in summary, our data demonstrates that UVB irradiation of murine skin drives IFNβ and ISG expression both locally and systemically in a STING-dependent manner, and that the absence of *Trim21* results in exacerbated responses both locally and systemically. The reduction of TRIM21 expression in SLE patient whole blood samples and association with cutaneous involvement strengthens our overall hypothesis that TRIM21 acts to restrain systemic IFN responses following environmental triggers such as UVB exposure of skin (as outlined in Figure 5).

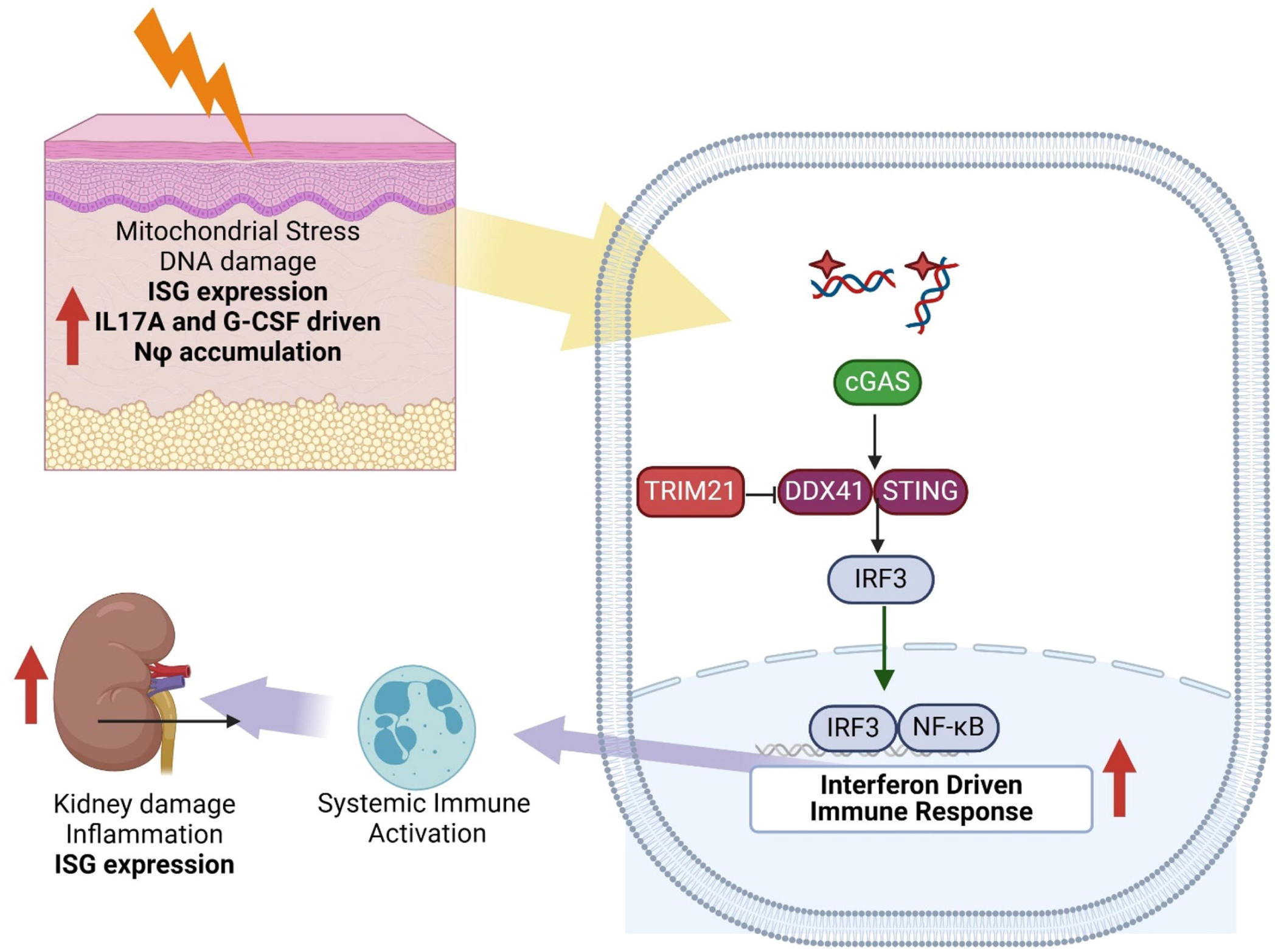

## Discussion

TRIM21 is an important autoantigen associated with cutaneous SLE and is also a critical negative regulator of type I IFN induction in response to RNA and DNA sensing. UVB is an important environmental trigger for both cutaneous SLE and in driving systemic flares, and has recently been shown in mice to activate both local and systemic type I IFN responses via a cGAS-STING dependent pathway. As loss of *Trim21* has previously been shown to result in skin inflammation we hypothesized that mice deficient in *Trim21* may be hypersensitive to UVB exposure. In keeping with this, we observed that *TRIM21*-deficient mice developed splenomegaly and systemic induction of type I IFNs in response to chronic UVB exposure compared to WT mice. This was accompanied by increased numbers of neutrophils and inflammatory monocytes, both in the circulation and in the spleen, and that this corresponded with increased expression of CCR2, a chemokine receptor critical for homing to inflamed tissues (such as the kidney or spleen). In addition, central memory CD4+ T cells were also increased in Trim21-deficient mice exposed to chronic low doses of UVB, suggesting that loss of *Trim21* results in enhanced CD4 T cell activation. Assessing potential targets for TRIM21 in this model revealed that the DNA sensor DDX41 and STING protein levels were stable in *Trim21^-/-^*murine cells, whereas both were decreased in a time-dependent manner in response to UVB or ISD transfection in cells derived from WT mice. Importantly, enhanced expression of *Ifn*β in *Trim21^-/-^* MDFs and BMDMs could be rescued by inhibiting signaling through STING, either through loss of *Tmem173* or using a STING-specific inhibitor. Importantly we observe decreased expression of TRIM21 in whole blood samples from SLE patients with cutaneous involvement, in addition to decreased expression of TRIM21 in the lesional skin of patients with cutaneous SLE compared to non-lesional skin. Our data underlines the importance of TRIM21 as a negative regulator of type I IFN responses and suggests that in the context of UVB-driven responses in SLE, loss of TRIM21 expression or activity may promote UVB-induced SLE exacerbations.

UV irradiation causes damage in cells via direct DNA damage leading to formation of pyrimidine dimers and activation of the DNA damage response via p53 activation. Indirect DNA damage results from oxidative stress and production of reactive oxygen species (ROS), driving oxidation of bases such as 8-oxo-2’-deoxyguanoisine (8-OH-dG) and single-stranded DNA breaks (16,17 of review). Indeed, we have previously shown that loss of Ogg1 in mice, an enzyme that degrades 8-OH-dG, develops a lupus-like disease and enhanced responses to pristane induced SLE, accompanied by increased expression of type I IFNs (Tumurkhuu G et al., 2020). In SLE, UVB exposure is accompanied by type I IFN and CXCL10 induction and are important in the pathogenesis of cutaneous involvement in SLE (3,39,42). In our studies, absence of Trim21 results in exacerbated IFNβ and CXCL10 in the skin following UVB exposure, indicating that Trim21 negatively regulates the production of these cytokines. In addition, we observe enhanced recruitment of CD45+ immune cells, and more importantly enhanced levels of expression of Siglec1, an IFN stimulated gene, on CD11b+ cells, indicating enhanced exposure of these cells to type I IFN.

The role of type I IFNs in cutaneous manifestations of SLE is well established, with production of type I IFN by epidermal cells and Langerhans cells responsible for priming inflammatory and apoptotic responses in the skin (2, 7, 38, 39). Recent single cell transcriptomic profiling of cells from lesional and non-lesional skin from SLE patients with active cutaneous involvement identified an overall high IFN-rich signature in both lesional and non-lesional skin, suggesting that normal appearing skin in SLE patients is actually primed immunologically and exists in a prelesional state (6). Another important implication from this analysis was that non classical monocytes in the periphery may be the source of the abundant levels of CD16^+^ dendritic cells observed in both lesional and non-lesional skin. These findings suggest that changes in the skin microenvironment promote immune cell infiltration and skewing towards pathogenic subtypes. In line with this, recent murine studies have shown that the transition from skin to systemic inflammation requires cGAS-STING signaling and that neutrophil and monocyte recruitment to inflamed skin in UVB models requires IFN signaling. Indeed, in the absence of cGAS the monocyte chemoattractant CCL2 (MCP-1) is diminished (36). Thus, in early responses to UVB, both CCL2 and IFN are required for neutrophil and monocyte recruitment to the skin. In our study we observe that systemic effects in response to skin exposure to UVB are exacerbated in *Trim21^-/-^*mice and that increased expression of Siglec1, an IFN-inducible gene, is observed on circulating CD11b^+^ cells. In addition, inflammatory CD11b^+^Ly6C^+^ monocytes show increased expression of the chemokine receptor *Ccr2*, the receptor for CCL2, which is required to home cells to inflamed tissues, including the skin and kidneys (1, 21, 35, 36, 40).

UVB also induces rapid accumulation of neutrophils to the skin in murine models. In the absence of *Trim21*, no differences in neutrophil numbers in the skin following UVB exposure compared to WT are observed. However, an increase in both relative percentage and absolute numbers of neutrophils in the blood and spleen of UVB-exposed *Trim21^-/-^*mice were seen. This is supported by increased expression of genes associated with neutrophil degranulation and myeloid cell activation and differentiation in *Trim21^-/-^*splenocytes, suggesting potentially that UVB exposure triggers emergency granulopoiesis in the *Trim21^-/-^* mice. This is in keeping with a recent study that examined the kinetics of neutrophil recruitment to the skin in C57BL/6 mice exposed acutely to UVB (35). In this study the authors observed that neutrophils trafficked to the skin within 24 hours post-UVB exposure and remained there for up to 6 days. This was accompanied by an increase in neutrophils systemically and in the spleen, in accordance with our own findings.

While TRIM21 is an important autoantigen in SLE and Sjogren’s disease, its role in the pathobiology of SLE and Sjogren’s has been somewhat ambiguous. As an autoantigen it has been shown to be extruded on blebs of apoptosing keratinocytes in response to UVB irradiation – thus promoting autoantibody formation. It is also an IFN regulated gene, thus its expression is frequently increased in SLE peripheral immune cells. However, the anomaly of enhanced TRIM21 expression and yet decreased regulation of DNA sensing in SLE immune cells has not been explained to date. In this study however we observe that decreased *TRIM21* expression is associated with cutaneous involvement in SLE patients, in keeping with loss of regulation of IFNb-dependent pathways. However, an in depth analysis of TRIM21 expression and function in SLE patient immune cells and stromal cells is required to fully understand how TRIM21 contributes to autoimmune pathology in the context of SLE. In summary we have identified a role for TRIM21 as a gatekeeper against UVB-induced systemic disease in mice (as shown in Figure 5), via its ability to regulate cGAS-STING dependent signaling in immune cells and stromal cells both at the local and systemic level.

