## Supplementary figures and images for "Loss of TRIM21 drives UVB-induced systemic inflammation by regulating DNA-sensing pathways"

### Supplemental Figures

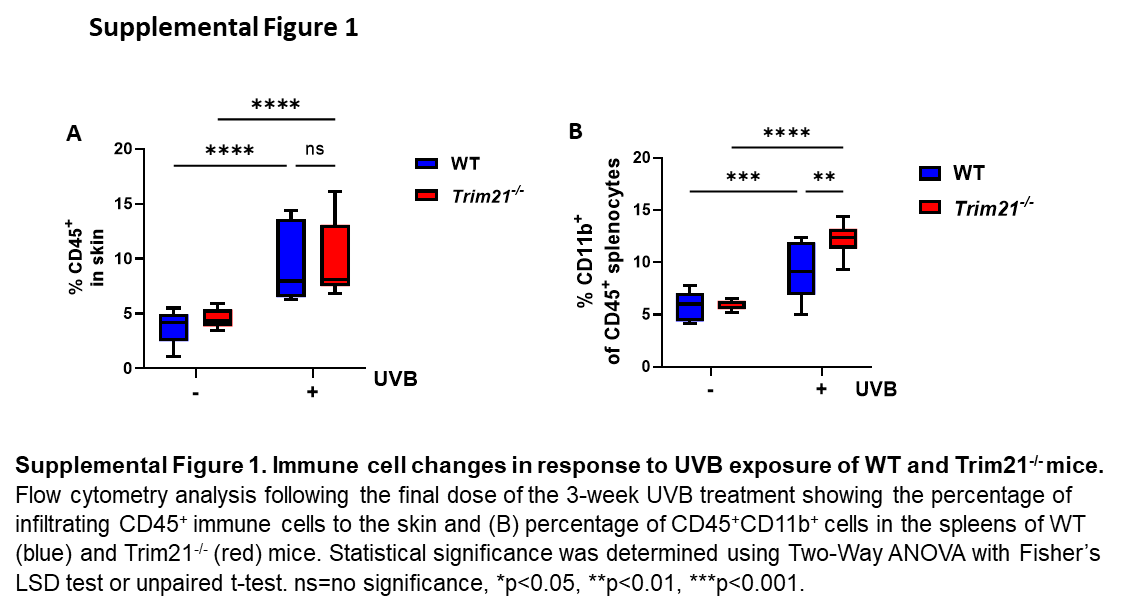

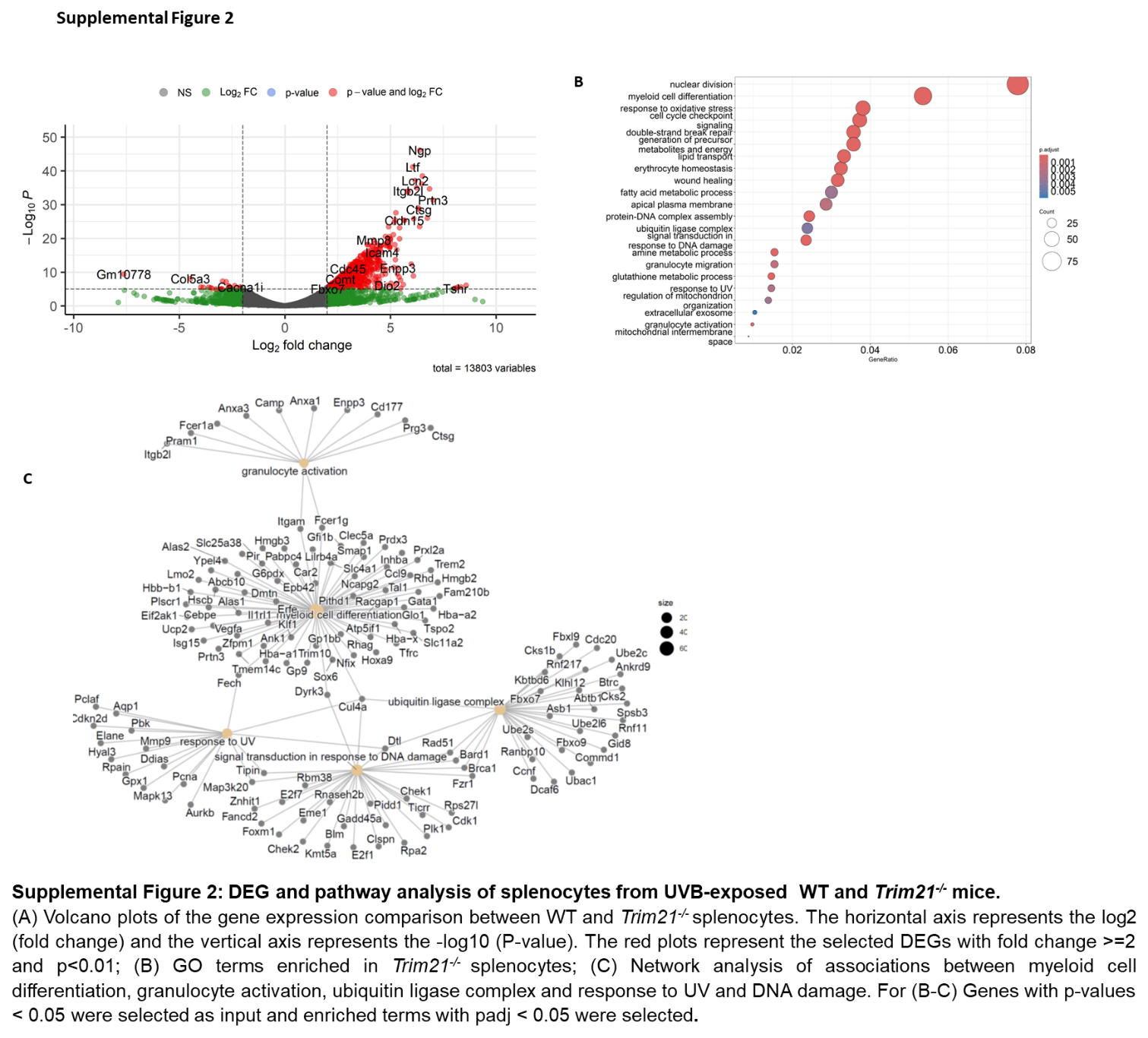

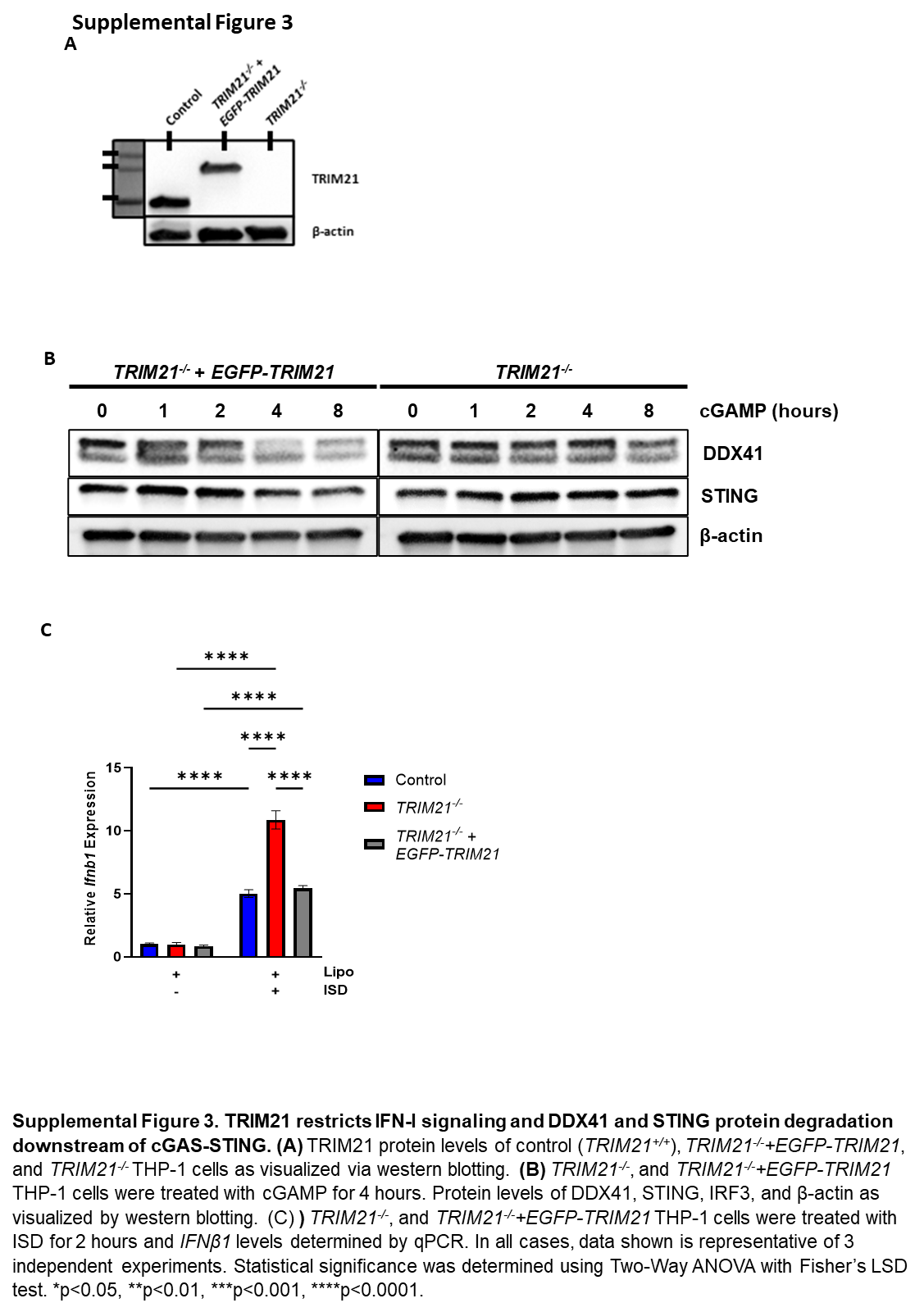

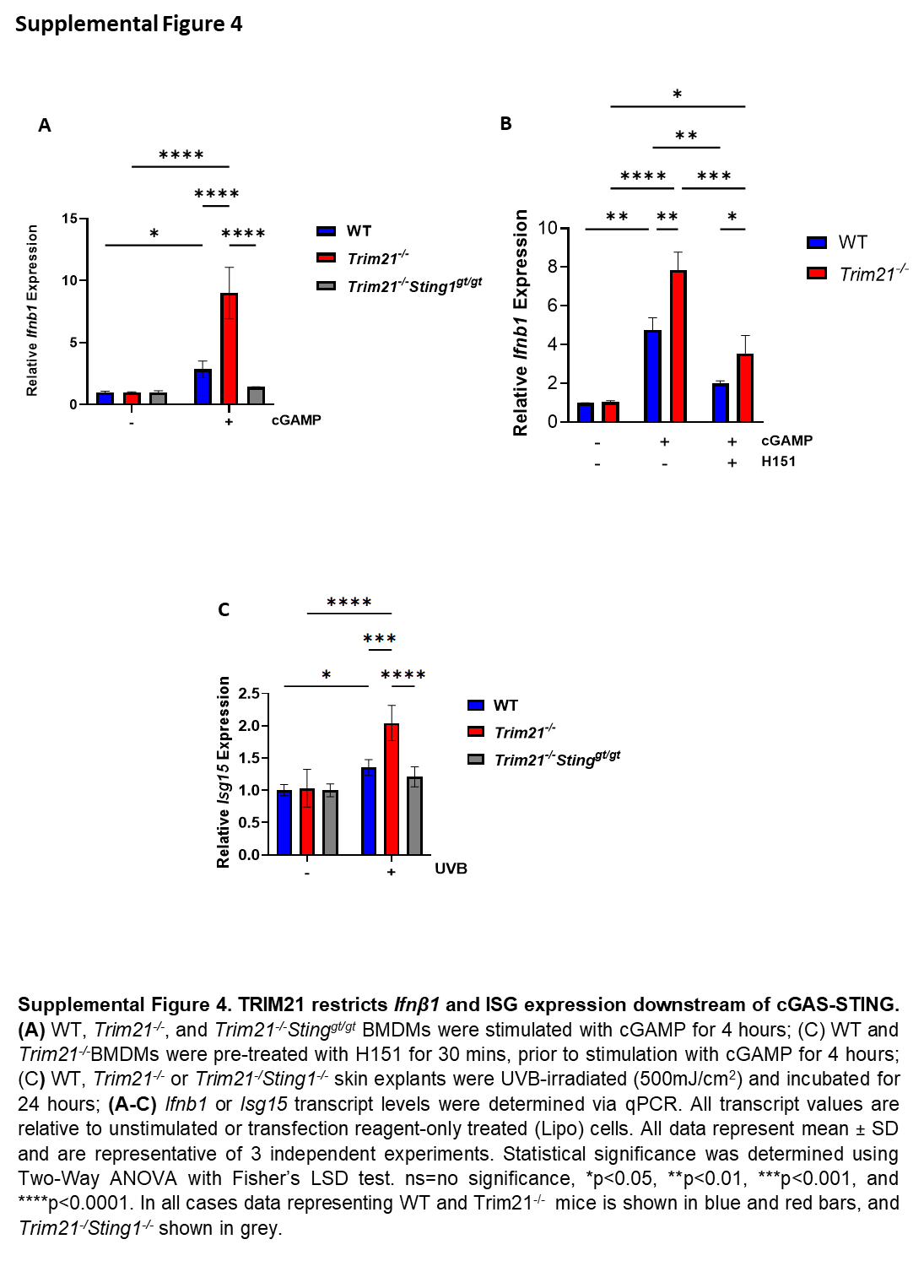
